# Biochar suppression of antibiotic resistance genes in the soil microbiome

**DOI:** 10.64898/2026.09.11.750991

**Authors:** Thomas Brook, Deepal Bhatt, Brian Reid, Marcela Hernández, Melisa Olivelli

## Abstract

Antimicrobial resistance (AMR) in soils frequently arises from low-level antibiotic presence in agriculture systems where livestock receive antibiotics for disease prevention. Here we assess the influence of oak (*Quercus robur*) wood biochar (pyrolysis at 760°C) on the microbiome of a parkland soil, after 300 days of incubation, in the presence and absence of the antibiotic sulphamethazine (SMZ). Stable isotope probing coupled to metagenomics was carried out to identify metabolically active microorganisms and quantify antimicrobial resistance gene (ARG) prevalence. Across all treatments, Pseudomonadota and Actinobacteriota dominated (63–78%), with Pseudomonadota enriched in the biologically active heavy H_2_^18^O fraction. Gemmatimonadota increased, and Bacillota decreased in heavy fractions, while minor phyla displayed variable shifts. Biochar significantly reduced ARG prevalence in the active heavy H_2_^18^O metagenome from 23,485 ARGs in control soils to 10,002 ARGs in biochar-amended soils, while SMZ alone increased to 26,193 ARGs found in their metagenome. Combined biochar and SMZ treatment resulted in lower ARGs (16,890 ARGs) than both control and SMZ-only soils, likely due to a combination of ARG adsorption, antibiotic immobilisation, and microbial community restructuring. These results highlight the potential of biochar as a soil amendment for mitigating the proliferation of AMR in agriculture environments.

## 1. Introduction

Soil is an important reservoir of antimicrobial resistance genes (ARGs), with studies identifying approximately 30% of all known ARGs in sequence databases to be found within soil (Nesme et al., 2014). Despite the historic presence of resistance genes, the human use of antibiotics has put a much higher selective pressure on bacteria to achieve resistance.

This prevalence of antimicrobial resistance (AMR) is a major threat to the efficacy of modern medicine. Awareness of this issue has pushed clinicians to limit prescription of antibiotics and better educate patients on the correct way to complete their antibiotic treatment courses to reduce the selective pressure for resistance in clinical settings (Allen et al., 2010). In contrast, in several countries, the agricultural industry continues to use antibiotics as prophylactics in animal food and water supplies – even in the absence of infection – to help promote growth and prevent common infections (Liyanage and Pathmalal, 2017).

Compounding the problem is often poorly absorption of active compounds, resulting in excretion rates of between 30-90% (Sarmah et al., 2006). This high potential for active compounds within animal excrement, sustains a potential to transfer ARGs and antibiotics into soil (Allen et al., 2010; Thanner *et al*., 2016; Chen et al., 2019).

Given the environmental risks posed by veterinary antibiotics, strategies to mitigate their persistence and impact in agricultural soils have become a global priority. Recent attention has turned to sustainability mitigation strategies, to reduce antibiotic microbial bioavailability, their selective pressure on soil microbiomes, and the propagation of AMR/ARGs (Fang et al., 2023; Olanrewaju and Bezuidenhout, 2025; Idress et al., 2026). A candidate material in this regard is biochar; a carbon-rich solid produced when organic matter is thermally decomposed under oxygen limited pyrolysis conditions (Freddo et al., 2012). Owing to large surface areas and reactive functional groups, biochar has been shown to be effective in the adsorption of a wide variety of contaminants (Khan et al., 2013, 2014; Waqas et al., 2016; Zama et al., 2017, 2018), including antibiotics (Krasucka et al., 2021). The adsorption of antibiotics to the surface of biochar results in the limitation of their interaction with the surrounding soil and its microbial community, thereby lowering the selective pressure towards AMR, and the spread of ARGs (Du et al., 2023). In addition, biochar has numerous beneficial properties when incorporated into soils; for example, it has been shown to increase soil fertility, increase soil microbial activity, increase water retention, and limit nutrient leeching (Krasucka et al., 2021; Du et al., 2023).

To accurately assess the prevalence and proliferation of ARGs within the soil microbiome – particularly in response to antibiotic and biochar exposure – it is essential to be able to distinguish between DNA derived from actively reproducing bacterial species from that of extracellular DNA (eDNA) originated from dead or inactive cells, which can persist in soils at high abundance (Chen et al., 2019). Previous studies have shown inconsistent results, ranging from no significant change in the soil microbiome to complete alteration upon exposure to antibiotics (Urra et al., 2019; Zheng et al., 2021). This variability has been partially attributed to the inclusion of eDNA in analyses, which can obscure signals from metabolically active populations. To address this limitation, we employed stable isotope probing (SIP) with H_2_^18^O to selectively label DNA from active microorganisms. Because water is required for cellular growth and maintenance, incorporation of ^18^O into newly synthesised DNA enables the separation of active and inactive fractions based on density. This approach provides a robust means of identifying metabolically active members of the soil microbiome, with more active organisms incorporating the label more rapidly, while all active taxa are expected to become labelled over extended incubation periods (Hernández et al., 2023).

The aim of our research was to evaluate the influence of biochar on the microbiome of a parkland soil and to evaluate the influence of biochar on ARG proliferation in the presence and absence of the veterinary drug sulfamethazine (SMZ). We selected SMZ as it is widely used in agriculture for the treatment for bacterial diphtheria, pneumonia and scours, in addition to bovine respiratory disease and necrotic pododermatitis (Sarmah et al., 2006).

Further, in a study covering six US states, SMZ has been reported to be the second most frequently detected antibiotic in swine and poultry liquid waste (Meyer et al., 2003). In order to target the biologically active fraction of the soil metagenome, we used SIP with H_2_^18^O.

## 2. Material and Methods

### 2.1. Chemicals

Caesium Chloride and H_2_^18^O, 97 atom (%) were purchased from Merck (Gillingham, UK). Sulfamethazine (99%) was purchased from Fischer Scientific. Biochar was synthesized using oak (*Quercus robur*) wood chips these were loaded into a cylindrical pyrolysis chamber (Φ150mm x 600mm). Pyrolysis occurred at 760°C as a downward moving gasification propagated through the fixed bed of wood chips. Resultant biochar was crushed until it passed through a 2mm sieve.

### 2.2. Soil incubations and stable isotope probing

Parkland soil, with low possible of prior antibiotic exposure, was sampled on the grounds of the University of East Anglia campus. Soil had a sandy loam texture with low organic matter content (2.6%) and pH (6.8). Soil was sieved (2mm) and microcosms prepared: soil only (control), soil with SMZ (50 ppm), soil with biochar (0.5% (w/w)), and soil with biochar and SMZ (50 ppm and 0.5% (w/w)). Each microcosm was prepared in triplicate and incubated for 300 d before six subsamples (1g) were taken from each microcosm. Samples (in triplicate) were incubated for 48 h with either (0.2ml) of H_2_^18^O or H₂¹⁶O (control), enabling incorporation of ¹⁸O into newly synthesised DNA of actively growing microorganisms. Following incubation, DNA was extracted and subjected to caesium chloride density gradient ultracentrifugation (45000 rpm for 44 hours), allowing separation of isotopically labelled (“heavy”) and unlabelled (“light”) DNA fractions based on buoyant density (Neufeld et al., 2007; Jia et al., 2019). The buoyant density of each fraction was calculated by the following empirical formula: ρ = -75.9318 + 99.2031*x* - 31.2551*x*^2^, where ρ denotes buoyant density (g/ml) and *x* denotes refractive index (Jia et al., 2019). In total, 12 fractions were collected in each gradient and DNA was purified via polyethylene glycol (PEG 6000) precipitation. Purified DNA was stored at −20 °C prior to amplicon and shotgun sequencing performed by Novogene (UK).

### 2.3. Total microbial community composition

The Novogene results for the 16S rRNA gene include Operational Taxonomic Unit (OTU) tables generated using the Silva 138.1 database (Quast et al., 2012). Relative abundance analysis of the 16S rRNA gene OTUs was completed in RStudio (v2024.09.0+375). For each soil treatment sample, OTUs were filtered to those of the bacterial kingdom, aggregate absolute counts for each phylum were calculated, and the percentage abundance of each phylum determined within the samples. Each treatment was then plotted using ggplot2 (v3.5.1), faceting the samples by whether they were incubated in H_2_^16^O or H_2_^18^O, then by light or heavy DNA-SIP fractions. Statistical analysis of the relative abundance of the microbial communities within the soil microbiomes was compared using the corncob taxon regression model available in R (Martin et al., 2020; Martin, 2024). For each soil treatment, the light H_2_^16^O fraction was used as reference and compared to the abundance levels of heavy H_2_^16^O, light H_2_^18^O and heavy H_2_^18^O fractions. The analysis was performed using a false discovery rate threshold of 5%. Alpha diversity indices measurements calculated from absolute OTU counts from Novogene using the R package vegan (v2.7-2). Simpson’s Index was calculated as Simpson’s Index of Diversity (1-D).

### 2.4. ARGs abundance

For the metagenomic data, we followed a similar bioinformatic pipeline previously described (Hernández et al., 2020). The raw read information was assessed for quality, ensuring no low-quality read information was present. This was achieved using FastQC (v0.11.8) with default parameters (Andrews, 2010). High quality reads were then assembled into de novo scaffolds utilising the genome assembler SPAdes (v3.14.0) (Prjibelski et al., 2020), with the parameters: meta, only-assembler and phred-offset 33. This ensured only assembly was performed, for a metagenomic sample with a PHRED quality offset of 33. The sequence aligner tool BBMap (v38.86) (Bushnell, 2014) and its associated K-mer based sequence trimming tool BBDuk, were used to trim the SPAdes scaffolds to those that were ≥ 1 kilobases (kb) in length using default parameters, and a minlen parameter of 1000. The scaffolds were then binned using the metaWRAP (v1.2.1) (Uritskiy et al., 2018) binning module, specifying for binning with metaBAT, MaxBin, and CONCOCT simultaneously.

To get an estimate of the total ARG abundance present within each soil microbiome, resistance gene identifier (RGI) analysis was performed on the unbinned ≥ 1kb scaffolds for each treatment. This was performed using the comprehensive antibiotic resistance database (CARD) (McArthur et al., 2013; Alcock et al., 2020), and its RGI tool (v6.0.5), which can perform read-based annotations of identified ARGs. The results were further analysed by identifying the total ARG matches, those which were strict matches, and those that were linked to sulphonamide resistance. RGI uses a Protein Homolog Model (PHM) to detect protein sequences based on their similarity to a curated reference sequence, using the CARD curated BLASTP bitscore cut-off. A perfect RGI match is 100% identical to the reference protein sequence across its entire length. A Strict RGI match meets or exceeds the BLASTP bit-score cut-off, whereas a Loose RGI match falls below the BLASTP bit-score cut-off (https://card.mcmaster.ca/ontology/40292). RGI was applied to scaffolds >1 kb from each metagenome.

## 3. Results

### 3.1. Relative abundance and corncob analysis of 16S rRNA gene

16S rRNA gene were used to produce relative abundance. The four treatment types were included: unaltered soil, soil with biochar, soil with SMZ, and soil with both SMZ and biochar. For each treatment type, results were faceted by water type (H_2_^16^O or H_2_^18^O) used during incubation, and whether the fractions were designated light or heavy during DNA-SIP fractionation. The top two most abundant phyla were consistently either *Pseudomonadota* or *Actinomycetota* across all treatments and represent between 63 to 78% of the total abundance regardless of treatment or DNA-SIP fraction (Fig. 1). However, the most abundant phyla typically switch from *Actinomycetota* to *Pseudomonadota* when analysing the biologically active heavy H_2_^18^O fraction compared to the other fractions. All soil treatments saw an increase in *Pseudomonadota* and reduction *Actinomycetota,* when comparing their light H_2_^16^O and heavy H_2_^18^O fractions. This is most apparent in the biochar treatment where a 15% increase in abundance of *Pseudomonadota* and a 15% reduction of *Actinomycetota* was observed (Fig. 1B). Similarly, the third and fourth most abundant phyla were typically *Gemmatimonadota* and *Bacillota* across all samples. *Gemmatimonadota* showed a consistently higher abundance in the heavy H_2_^18^O fraction (9.3-15.1%) compared to the other three fractions (5.5-10.5%). *Bacillota,* in contrast, was observed with a consistent reduction in abundance in the heavy H_2_^18^O fraction (1.3-3.6%) compared to other fractions (2.9-17.0%, Fig. 1). The remaining of the top 10 most abundant phyla (each individually represent between 0 to 6% of the relative abundance), consisted of *Acidobacteriota, Chloroflexota, Myxococcota, Planctomycetota, Bacteroidota,* and *Verrucomicrobiota* in descending abundance (average across all samples). The differences between these phyla when comparing to the heavy H_2_^18^O fraction varied in whether a decrease or increase was observed. There was a consistent reduction in *Acidobacteriota* (0.4-2.0%) and *Chloroflexota* (0.8-3.3%) and an increase for *Bacteroidota* (0.7-4.43%) phyla in the heavy H_2_^18^O fraction. The biochar treatment showed the least variation between samples (Fig 1B) and the clearest contrast in abundance levels when comparing the heavy H_2_^18^O fraction with all other fractions (Fig 1B). In contrast, both treatments including SMZ showed a much higher deviation between repetitions, particularly in the *Bacillota* of the light H_2_^18^O fraction (Fig 1C, 1D).

**Figure 1:**
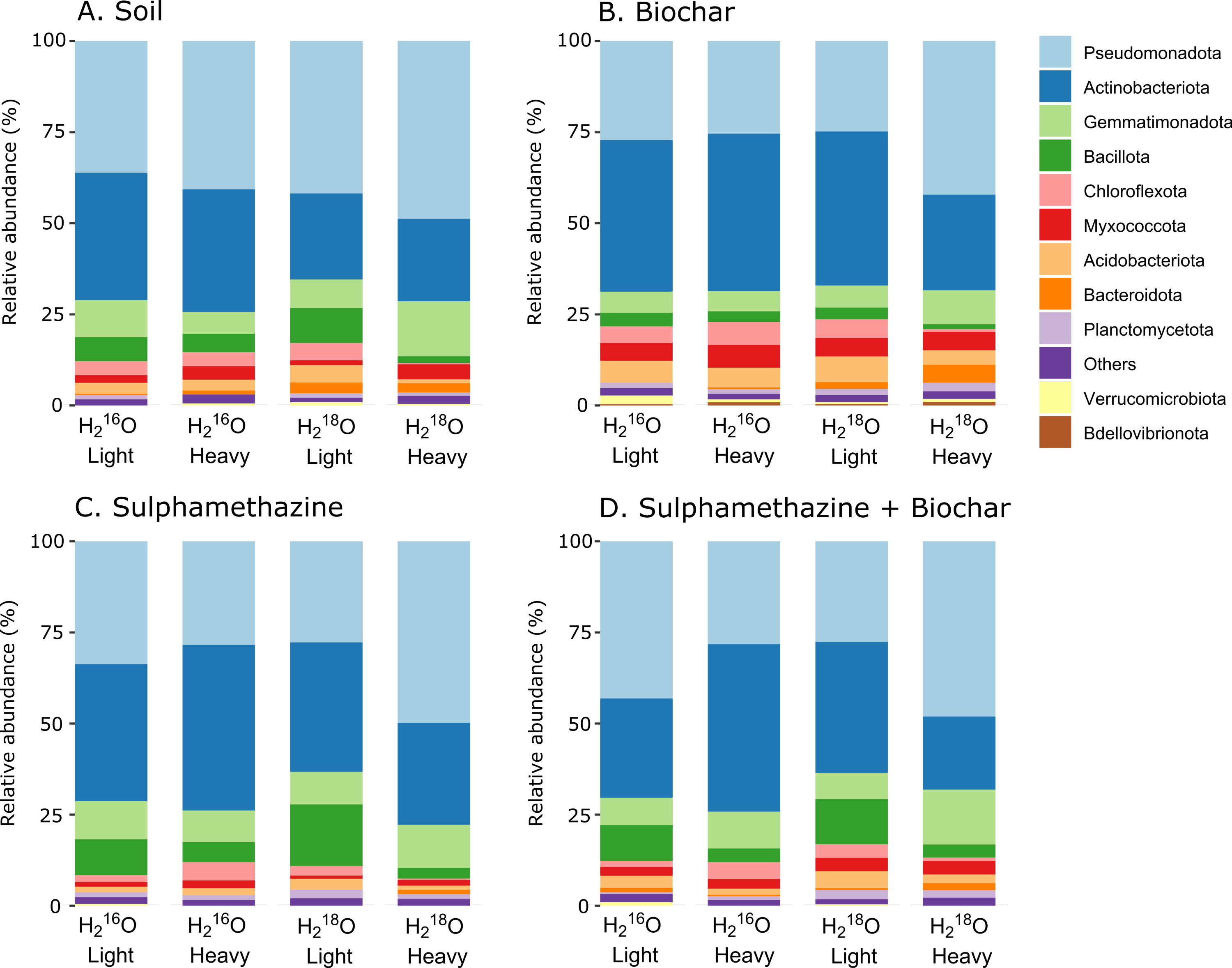
Mean relative abundance of microbial communities at the phylum level identified in the heavy and light DNA stable isotope probing fractions of 16S rRNA gene extracted from soils incubated with H_2_^16^O or H_2_^18^O, incubated under four treatment conditions: A. Unaltered soil; B. Soil with biochar; C. Soil with SMZ; D. Soil with SMZ and biochar. Mean values are calculated from triplicate samples.

To get further statistical comparisons with higher stringency between the metabolically active heavy H_2_^18^O fraction and the light fractions (not active), a taxonomic regression model (corncob) was applied to the OTU counts to determine differential abundance. Results include OTUs, which showed statistically significant differential abundance in the heavy H_2_^18^O, heavy H_2_^16^O, and light H_2_^18^O fractions, when compared to the light H_2_^16^O fraction, using a false discovery threshold of 5%. For all conditions, *Actinomycetota* OTUs was the most common phylum observed to reduce in abundance in the heavy H_2_^18^O fraction; ranging from 36 to 75 OTUs with significant reduction detected amongst the soil treatments (Fig. 2). The most common phylum that shows a significant increase, was from *Pseudomonadota* with 24 OTUs detected in the biochar treatment (Fig. 2B). This phylum does show ambiguity amongst OTUs, with both increases and decreases of OTU abundance seen in the same soil treatment. Whether higher counts of increases or decreases were detected, was dependent on soil treatment. The findings for *Gemmatimonadota* and *Bacillota* match those from the relative abundance results in that *Gemmatimonadota* OTUs were consistently observed at higher differential abundance whilst *Bacillota* were of lower abundance. Across all treatments, higher proportions of OTUs indicated significant reductions than increases. The lowest proportion was seen in the biochar treatment with 68% of OTUs showing a significant reduction (Fig. 2B), whilst 93% of OTUs with significant differential abundance in the SMZ treatment were reduced (Fig. 2C). *Bacillota* and *Chloroflexota* saw a consistent reduction in the heavy H_2_^18^O fraction, with only the SMZ with biochar treatment not exhibiting a significant reduction (Fig. 2). The biochar treatment resulted in the greatest disparity in the active microbiome, showing 205 OTUs and 9 phyla with significant differential abundance changes from the other fractions, in comparison to 125 and 3 for unaltered soil, 69 and 4 for SMZ and biochar regimes, and only 57 and 1 for SMZ regimes (Fig. 2).

**Figure 2:**
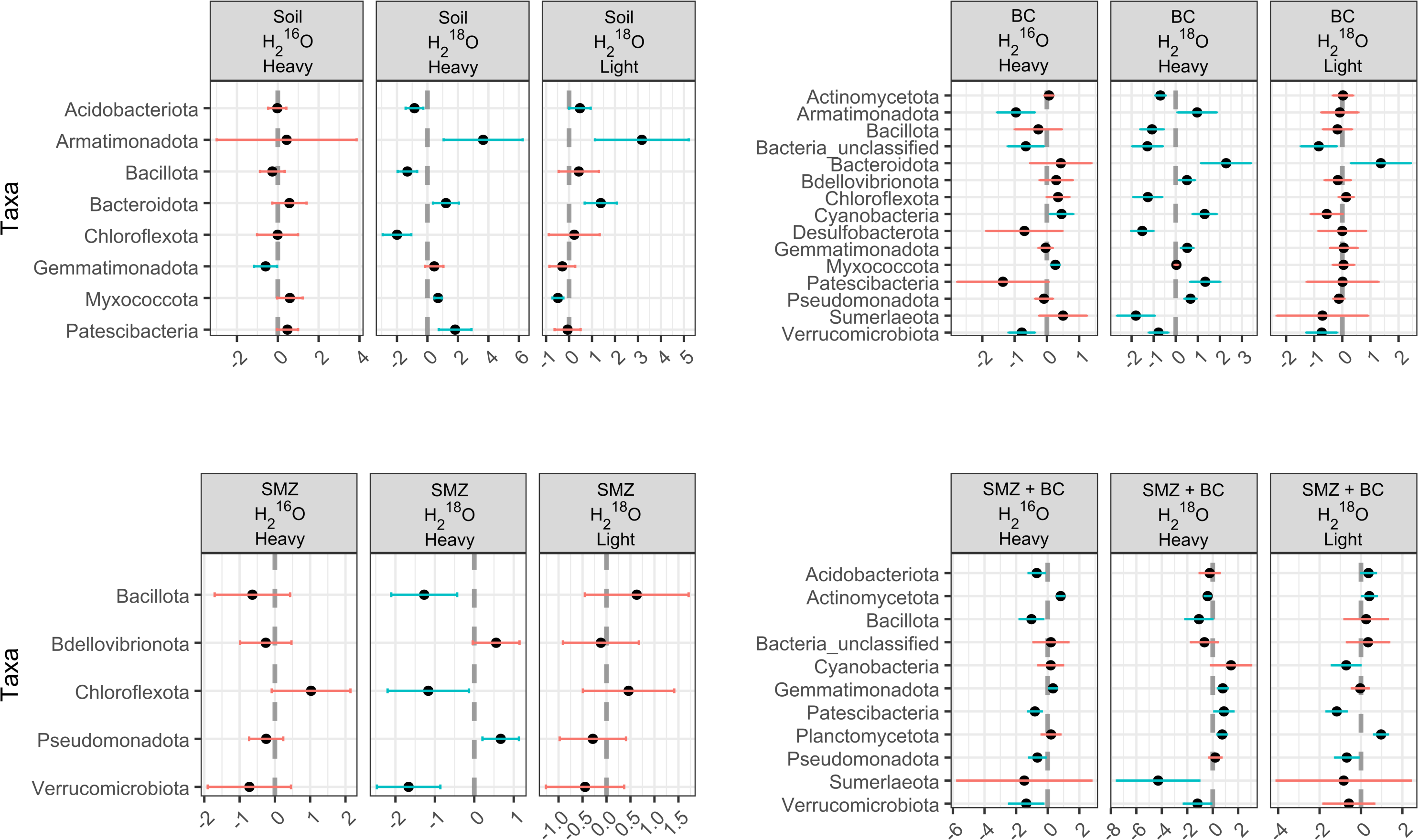
Corncob taxon regression analysis comparing the relative abundance of phyla when OTU abundance was aggregated by phyla. Using the light H_2_^16^O fraction as reference abundance and compared to the abundance levels of heavy H_2_^16^O, heavy H_2_^18^O and light H_2_^18^O fractions, using a false discovery rate threshold of 5%. Soil Treatments: Unaltered (soil), Biochar (BC), SMZ (Sulfamethazine), and SMZ and BC. Blue and red lines indicate true and false statistically significant difference, respectively.

At the class level, relative abundance within the heavy H_2_^18^O fraction revealed that the most abundant classes, based on the mean relative abundance across the four treatments, were Gammaproteobacteria (29.6%), Alphaproteobacteria (19.2%), Actinobacteria (15.9%), Gemmatimonadetes (12.7%), and Thermoleophilia (7.3%) (Fig. 3A). Biochar treated soil showed a lower proportion of Gammaproteobacteria, 19% compared to 31.5-35.1% in the other three soil treatments. Biochar also showed the highest proportion of Alphaproteobacteria compared to any other treatment, 23.0% in comparison to 16.0% in untreated soil. SMZ treated soil showed a higher proportion of Actinobacteria than any other soil treatment, 21.3% versus 13.0%-15.8% in the remaining three soil treatments.

**Figure 3:**
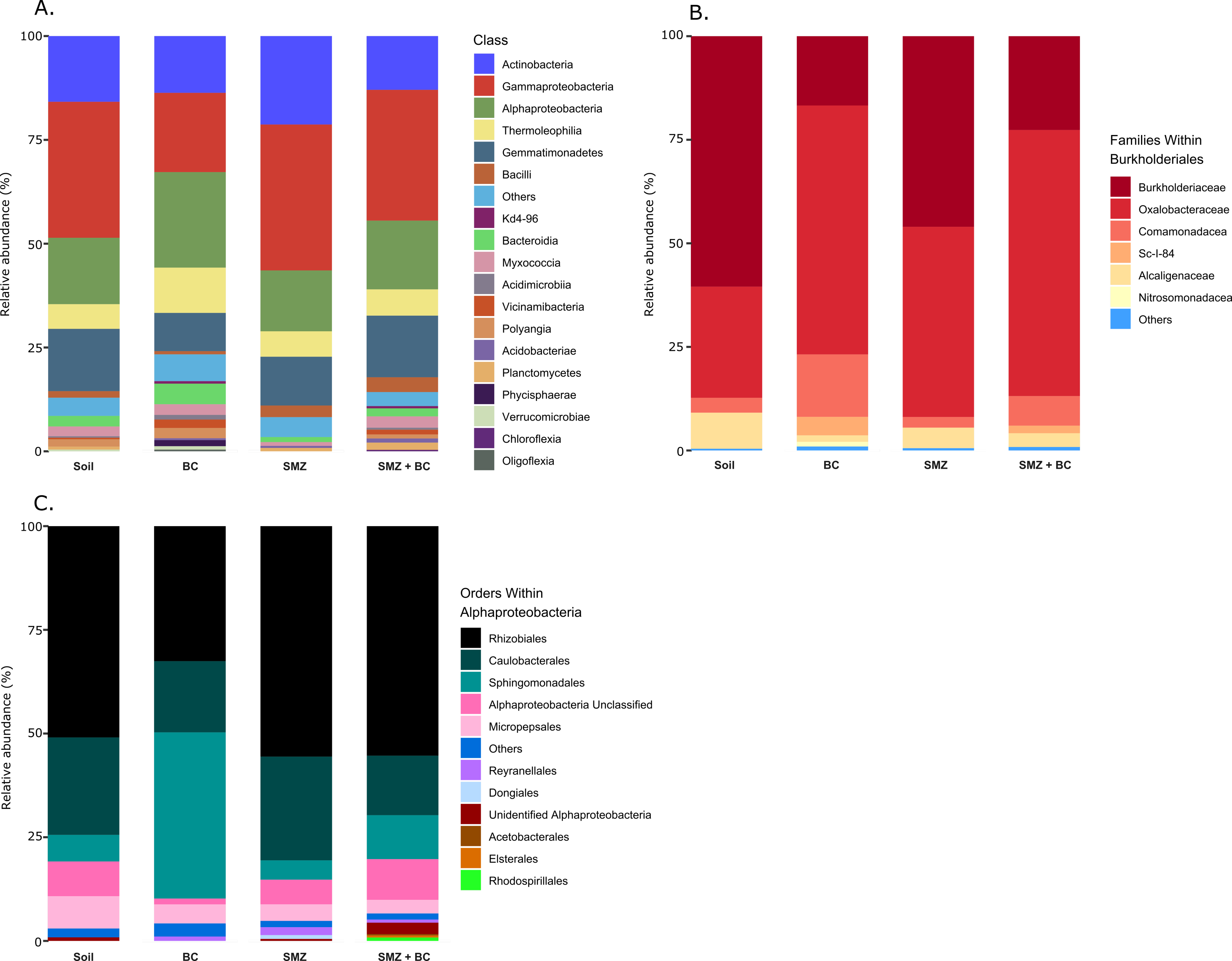
Mean relative abundance of microbial communities determined from 16S rRNA gene extractions from the heavy stable isotope probing fraction of soils incubated with H_2_^18^O under four treatment conditions: Unaltered soil; Soil with BC; Soil with SMZ; Soil with SMZ and BC. A. At the class taxonomic level; B. Filtered to the families within the order of Burholderiales; C. Filtered to the orders within the class Alphaproteobacteria. Mean values are calculated from triplicate samples.

Gemmatimonadetes showed similar proportions across all soils from 9.2% in biochar to 15.0% in untreated soil. Biochar also showed a higher proportion of Thermoleophilia, 10.9% in comparison to 5.9-6.3% in the other soil treatments. The class Gammaproteobacteria was predominately represented by the order Burkholderiales accross all soil treatments, ranging from 86-98% of sequences (Fig. 3B). The family Burkholderiaceae decreased from 60.5% in untreated soil to 16.8% in biochar-treated soils, whereas Oxalobacteraceae increased from 26.8% to 60.1%. In addition, the family SC-I-84 increased from <1% in untreated soils to 4.5% following biochar application.

Within the class Alphaproteobacteria (Fig. 3C), the order Rhizobiales decreased from 50.9% in untreated soil to 32.5% in biochar-treated soil, whereas Sphingomonadales increased from 6.4% to 40.0%. Within Sphingomonadaceae, the relative abundance of Porphyrobacter increased from 3.2% in untreated soil to 15.3% in SMZ-treated soils (Fig. S1). The taxon Plot4-2H12 also increased from 0.4% in untreated soil to 2.2% in SMZ-treated soil (Fig. S1). BC-treated soil showed a higher abundance of unclassified Sphingomonadaceae at 4.0% compared with < 1% in untreated soil (Fig. S1).

Biochar-treated soils within the heavy H_2_^18^O fractions showed greater microbial diversity compared to the other treatments (Fig. 4). Across all indices, biochar-treated soils exhibited the highest alpha diversity, whereas SMZ-treated soils showed the lowest. Soils treated with both SMZ and biochar consistently displayed lower diversity than untreated soils across all indices.

**Figure 4:**
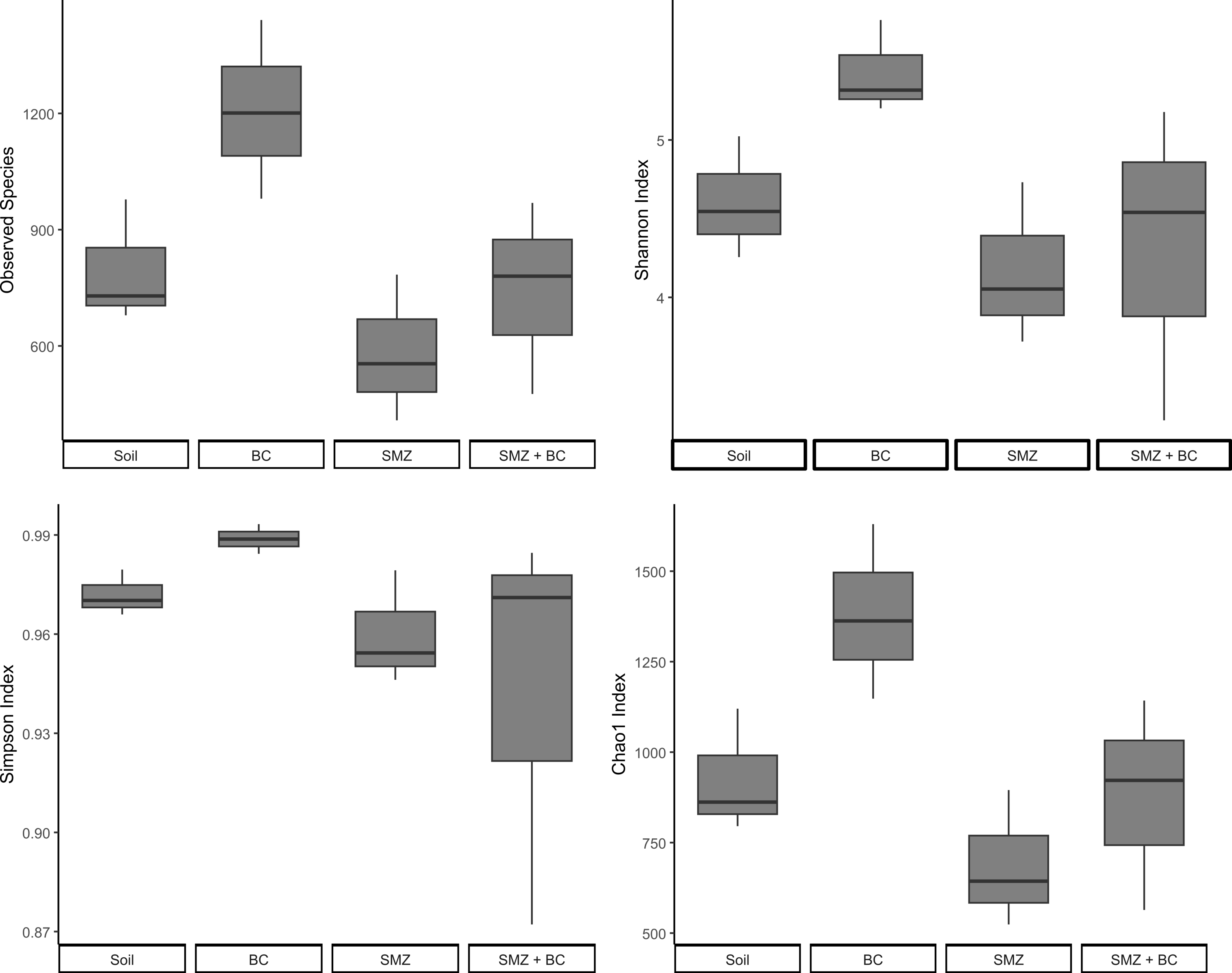
Alpha diversity of microbial communities in heavy fractions incubated with H_2_^18^O across all soil treatments. Alpha diversity was evaluated using observed species, the Shannon index, Simpson’s index, and the Chao1 index.

### 3.2. Metagenomic analysis and ARG abundance

The total ARG matches detected were identified using the default loose and strict matching criteria from RGI, with no perfect matches detected across in any metagenome. Biochar consistently showed the lowest abundance of ARGs across all three categories: total matches, strict matches and sulphonamide related ARGs, in both H_2_^18^O and H_2_^16^O heavy fractions, with 10002 and 9980 ARGs, respectively (Table 1). Across treatments, total ARG abundance increased in the following order: biochar, SMZ with biochar, Unaltered Soil, and SMZ, which showed the highest levels. This trend was consistent in both heavy fractions, with SMZ reaching 26193 ARGs in H_2_^18^O and 16150 in H_2_^16^O (Table 1). Biochar amendment substantially reduced ARG prevalence in the active metagenome, decreasing from 23,485 ARGs in control unaltered soils to 10,002 ARGs (Table 1). The combined biochar and SMZ treatment resulted in lower ARGs (16,890 ARGs) in H_2_^18^O than both unaltered soil and SMZ-only soils, likely due to a combination of ARG adsorption, antibiotic immobilisation, and shifts in microbial community structure.

**Table 1.**
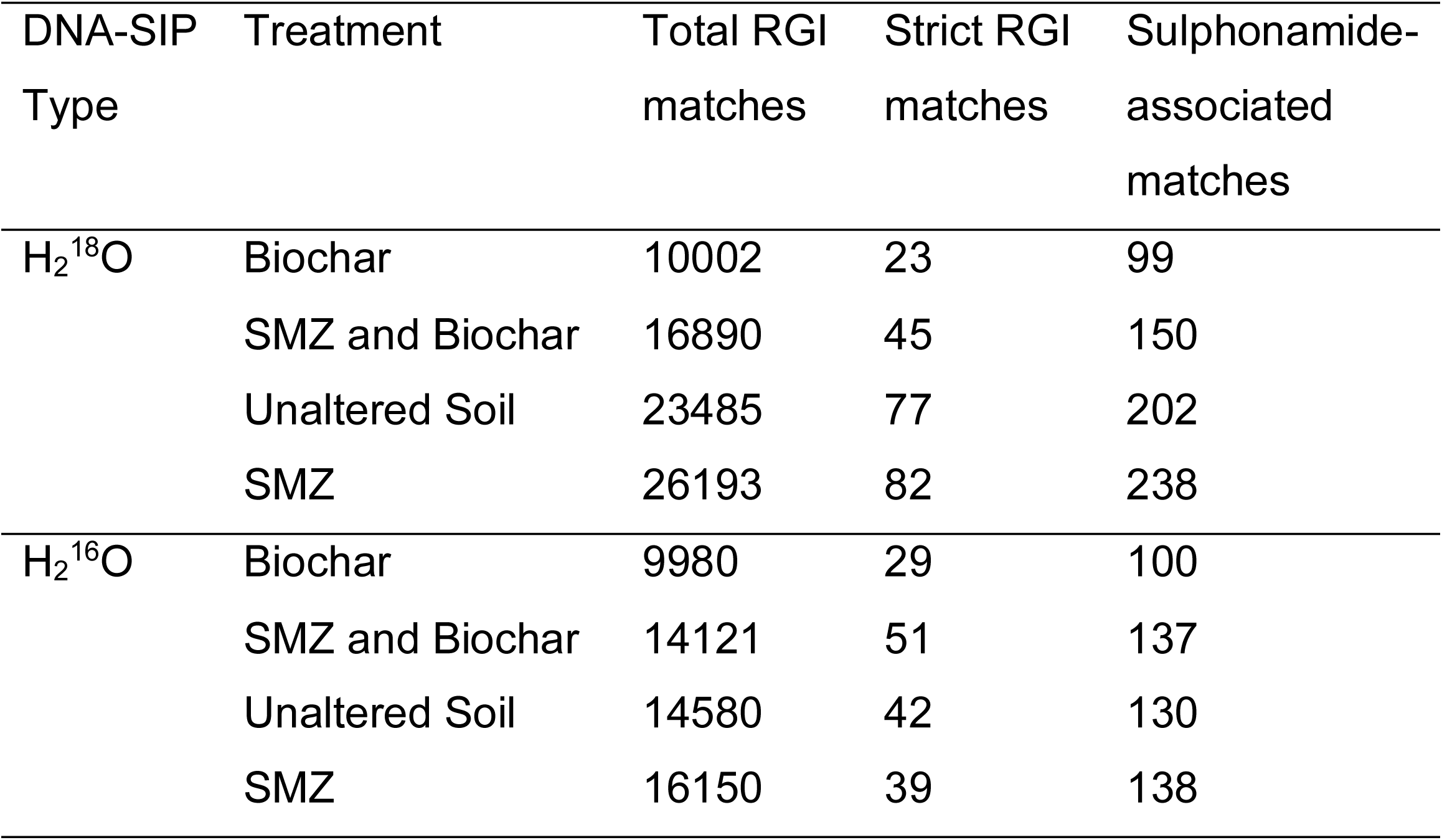
Summary of antibiotic resistance gene matches returned by the Resistance Gene Identifier (RGI) from the Comprehensive Antibiotic Resistance Database (CARD) from the unbinned ≥ 1 kilobase scaffolds of the metagenomic DNA samples.

| DNA-SIP Type | Treatment | Total RGI matches | Strict RGI matches | Sulphonamide-associated matches |
| --- | --- | --- | --- | --- |
| $\text{H}_2^{18}\text{O}$ | Biochar | 10002 | 23 | 99 |
|  | SMZ and Biochar | 16890 | 45 | 150 |
|  | Unaltered Soil | 23485 | 77 | 202 |
|  | SMZ | 26193 | 82 | 238 |
| $\text{H}_2^{16}\text{O}$ | Biochar | 9980 | 29 | 100 |
|  | SMZ and Biochar | 14121 | 51 | 137 |
|  | Unaltered Soil | 14580 | 42 | 130 |
|  | SMZ | 16150 | 39 | 138 |

## 4. Discussion

The relative abundance studies and corncob analyses highlight a clear separation between abundances seen in the active heavy H_2_^18^O fraction compared to all other DNA-SIP fractions. A high proportion of OTUs showed reduced abundance compared to an increase in the heavy H_2_^18^O fraction (data not shown). This pattern is consistent with the hypothesis that inactive microbial DNA or eDNA is primary present in H_2_^16^O DNA extractions. DNA extracted from the heavy H_2_^18^O fraction, incorporated the ^18^O provided during the incubation period, and must therefore have been recently metabolically active within the soil. Many OTUs that only have ^16^O present within DNA, such as in eDNA that is inactive, would be collected in the H_2_^16^O fractions, and the light H_2_^18^O fraction. These OTUs would have been filtered from the heavy H_2_^18^O fraction during DNA-SIP and, consequently, would show a significant reduction in abundance only in this fraction compared to all other fractions. We see evidence that this is the case, with 68-93% of OTUs with significant differential abundance, showing reduction in abundance across all treatments. We know from previous studies that ^18^O incorporation is stable and does not get easily exchanged with eDNA (Blazewicz and Schwartz, 2011), further supporting these findings. The results from our unaltered soil samples are perhaps the most conclusive evidence of this, in that the changes in OTUs abundances could not have been caused by any treatments, the only alterations are between the DNA-SIP incubation and fractionation. The phylum that is most frequently increased in the unaltered soil heavy H_2_^18^O fraction is *Pseudomonadota,* which is consistent with previous rewetting studies (Aanderud and Lennon, 2011); however, we see a far greater reduction in *Actinomycetota* than *Acidobacteriota* (Fig. 1). This suggests the changes in taxonomy are soil dependent, reinforcing the requirement of H_2_^18^O DNA-SIP incubation to accurately describe changes to the active microbiota in response to soil treatments. As such, many studies have included H_2_^18^O DNA-SIP in soil studies including rewetting (Aanderud and Lennon, 2011; Blazewicz et al., 2020), warming (Purcell et al., 2022) and fertilisation (Spohn et al., 2016).

Of all the treatments, only the biochar treatment showed a higher proportion of OTUs with increased abundance in the heavy H_2_^18^O fraction (Fig. 2). This may be attributed to the ability of biochar to reduce selection pressure from antibiotics present in the soil through adsorption (Pan, 2020). Consequently, OTUs lacking ARGs that confer protection against these antibiotics may have been able to replicate at higher frequencies. Alternatively, improvements in soil properties associated with biochar application, such as enhanced water retention, pH modifications, and increased nutrient availability (Li et al., 2019), may have favoured these OTUs. Additionally, biochar-treated soil showed the highest number of OTUs and phyla with significant differential abundance, suggesting that biochar has a stronger influence on microbiome taxonomic composition than SMZ. Evidence of this was also shown in the relative abundance analysis, which showed biochar treated soils diverged more from untreated soil than the SMZ treated soils (Fig. 3). The alpha diversity analysis (Fig. 4) supports these findings, with biochar treated soils showing higher levels of diversity across all indices measured. This increase is consistent with previous reports showing that biochar can enhance microbial diversity (Xiang et al., 2023; Xu et al., 2023), although the extent of this effect depends on soil quality (Li et al., 2020).

By analysing the abundance of ARGs within the unbinned metagenomic data for each treatment, we infer that the detected ARGs are present in proportions similar to those in the whole microbiome for each treatment. Under this assumption, ARG matches identified using RGI against the CARD database provide a basis for hypothesising the impact of biochar and SMZ on ARG prevalence in soils. Our primary focus is the biologically active microbial population, which is represented in the heavy DNA-SIP fraction derived from H_2_^18^O- incubated soil. We hypothesised that the introduction of SMZ would lead to a significant increase in ARG abundance within the active microbial metagenome compared with untreated soil, and that biochar application would limit or prevent this increase.

The results show that, although ARG abundance increased from untreated soil to SMZ-treated soil, the overall increase was relatively small across all categories (Table 1). The observed increase in sulphonamide-associated ARGs is consistent with the expectation that SMZ application promotes a higher prevalence of SMZ resistance ARGs. Notably, the co-application of biochar with SMZ restored ARG abundance to levels comparable with untreated soil and further reduced it to even lower proportions (Table 1). These results suggest that biochar not only mitigates the accumulation of ARGs induced by SMZ but also contributes to a broader reduction in ARG prevalence. This reduction may, in part, be attributed to the adsorption properties of biochar, which can limit interactions between soil microbes and antibiotics present in the environment (Pan, 2020). When applied in combination with SMZ; however, biochar adsorption sites may become saturated with the added antibiotic, as suggested by studies on adsorption equilibria (Du et al., 2023). This saturation could reduce the adsorption of other antibiotics, resulting in a higher prevalence of ARGs associated with those compounds.

In addition to the antibiotic adsorption, biochar has been shown to adsorb extracellular ARGs (eARGs), thereby preventing horizontal gene transfer (Fang et al., 2022; Wu et al., 2022). This mechanism could further reduce ARG propagation in biochar-treated soils, beyond those associated specifically with SMZ. For example, one study reported a >90% reduction in ARG abundance in wastewater when applying magnetically modified biochar (Fu et al., 2021). Our results suggest that the adsorption of eARGs by biochar may contribute more substantially to the observed reduction in ARGs abundance that antibiotic adsorption alone. This is supported by a similar study investigating the impact of SMZ and biochar in soil, which identified reductions in both eARGS and intracellular ARGs (iARGs) following biochar application (Qiu et al., 2021). Consistent with this, biochar-only treatment in our study resulted in the lowest ARG prevalence across all three categories (Table 1). Finally, the difference in total ARG counts between SMZ plus biochar, and biochar-only treatments was approximately 6900, whereas only 150 ARGs were detected with SMZ resistance (Table 1), further indicating that the observed effects extend beyond SMZ-specific resistance.

The results for the heavy fraction of the H_2_^16^O incubated soil show no clear pattern among the treatment types, although biochar application still consistently reduced ARG prevalence compared with untreated soil (Table 1). One potential explanation for these inconsistencies, relative to the H_2_^18^O results, is that the heavy fractions of the H_2_^16^O metagenome likely contains eDNA of dead or inactive bacteria commonly present in soil (Chen et al., 2019). As a result, it is not possible to determine what proportion of the metagenome was present prior to treatment and was therefore unaffected by the application of biochar or SMZ. Together with changes observed in the relative abundance of the heavy H_2_^18^O microbiome, these findings underscore the importance of confirming that observed shifts occur within the biologically active microbial population directly exposed to the treatments. These results may also help explain discrepancies reported in previous studies (Urra et al., 2019; Zheng et al., 2021) and supports the inclusion of H_2_^18^O DNA-SIP for future studies of metagenomic responses.

## 5. Conclusions

The divergence in taxonomic populations between the heavy H_2_^18^O DNA-SIP fractions and other fractions in our relative abundance analysis highlights the importance of identifying actively replicating DNA when assessing changes in soil metagenomes under changing conditions. This is further supported by the inconsistent ARG abundance patterns detected in the heavy H_2_^16^O DNA-SIP fractions compared to the heavy H_2_^18^O fraction, indicating that DNA extracted without isotopic labelling may not accurately represent the biologically active microbiome influences by experimental treatments. The observed reductions in ARG abundance, both in the presence and absence of SMZ, to levels below those in control soil provide strong evidence that biochar not only mitigates increases in ARG through antibiotic adsorption but may also reduce horizontal gene transfer by adsorbing eARGs. These findings are consistent with previous studies and demonstrate the potential of biochar to substantially reduce ARG prevalence in soil microbial communities. A key long-term goal is to evaluate the efficacy and persistence of biochar application in agricultural soils, particularly those exposed to high antibiotic loads, in limiting ARG dissemination and reducing the risk of transfer to human pathogens. Future studies should investigate whether biochar remains effective over extended periods and under repeated antibiotic exposure, and whether it represents a cost-effective strategy for sustainable ARG mitigation without requiring frequent reapplication.

## Acknowledgements

The authors would like to thank Dr. Shamik Roy for his assistance with bioinformatics, particularly with corncob.

## Data Availability

Sequencing data have been deposited in the NCBI Sequence Read Archive (SRA) under accession code PRJNA1469527 for both amplicon and metagenomic DNA sequencing.

## Author Contributions

MH: Supervision. MH, MO, and BR: Resources, Investigation, and Visualisation. MH, MO, and BR: conceived the ideas for the manuscript. MO: performed the initial incubations; DB: performed laboratory work, TB and MH: analysed the data. TB: Developed and designed the figures. TB and MH drafted the paper. All authors contributed to its revision and approved the final version.

