## Supplementary Information for "Biochar suppression of antibiotic resistance genes in the soil microbiome"

**This PDF file includes:**

Figure S1

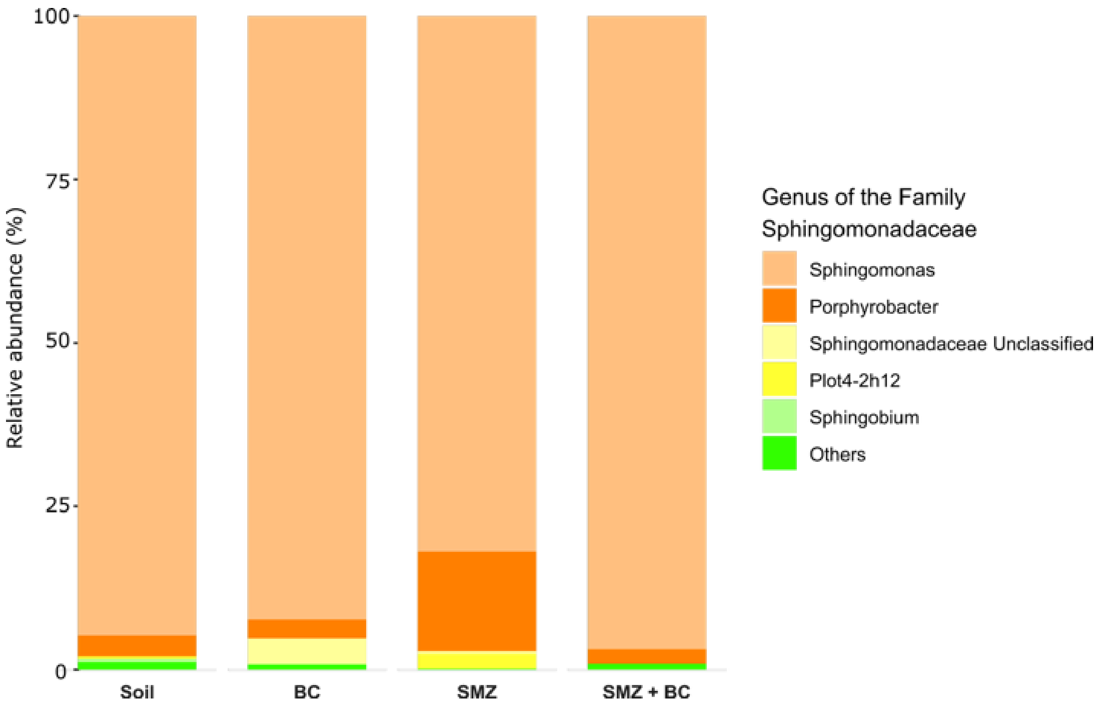


Figure S1: Mean relative abundance of genera within the family Sphingomonadaceae, determined from 16S rRNA gene extractions from the heavy stable isotope probing fraction of soils incubated with $H_2{}^{18}O$ under four treatment conditions: Unaltered soil; Soil with BC; Soil with SMZ; Soil with SMZ and BC. Mean values are calculated from triplicate samples.
